# Quantitative single-cell transcriptome-based ranking of engineered AAVs in human retinal explants

**DOI:** 10.1101/2021.09.27.461256

**Authors:** Zhouhuan Xi, Bilge E. Öztürk, Molly E. Johnson, William R. Stauffer, Leah C. Byrne

**Author notes:** These authors contributed equally to this work (Zhouhuan Xi, Bilge E. Öztürk). Correspondence should be addressed to: Leah Byrne,.

## Abstract

Gene therapy is a rapidly developing field, and adeno-associated virus (AAV) is a leading viral vector candidate for therapeutic gene delivery. Newly engineered AAVs with improved abilities are now entering the clinic. It has proven challenging, however, to predict the translational potential of gene therapies developed in animal models, due to cross-species differences. Human retinal explants are the only available model of fully developed human retinal tissue, and are thus important for the validation of candidate AAV vectors. In this study, we evaluated 18 wildtype and engineered AAV capsids in human retinal explants using a recently developed single-cell RNA-Seq AAV engineering pipeline (scAAVengr). Human retinal explants retained the same major cell types as fresh retina, with similar expression of cell-specific markers, except for a cone population with altered expression of cone-specific genes. The efficiency and tropism of AAVs in human explants were quantified, with single-cell resolution. The top performing serotypes, K91, K912, and 7m8, were further validated in non-human primate and human retinal explants. Together, this study provides detailed information about the transcriptome profiles of retinal explants, and quantifies the infectivity of leading AAV serotypes in human retina, accelerating the translation of retinal gene therapies to the clinic.

**Graphic abstract:** 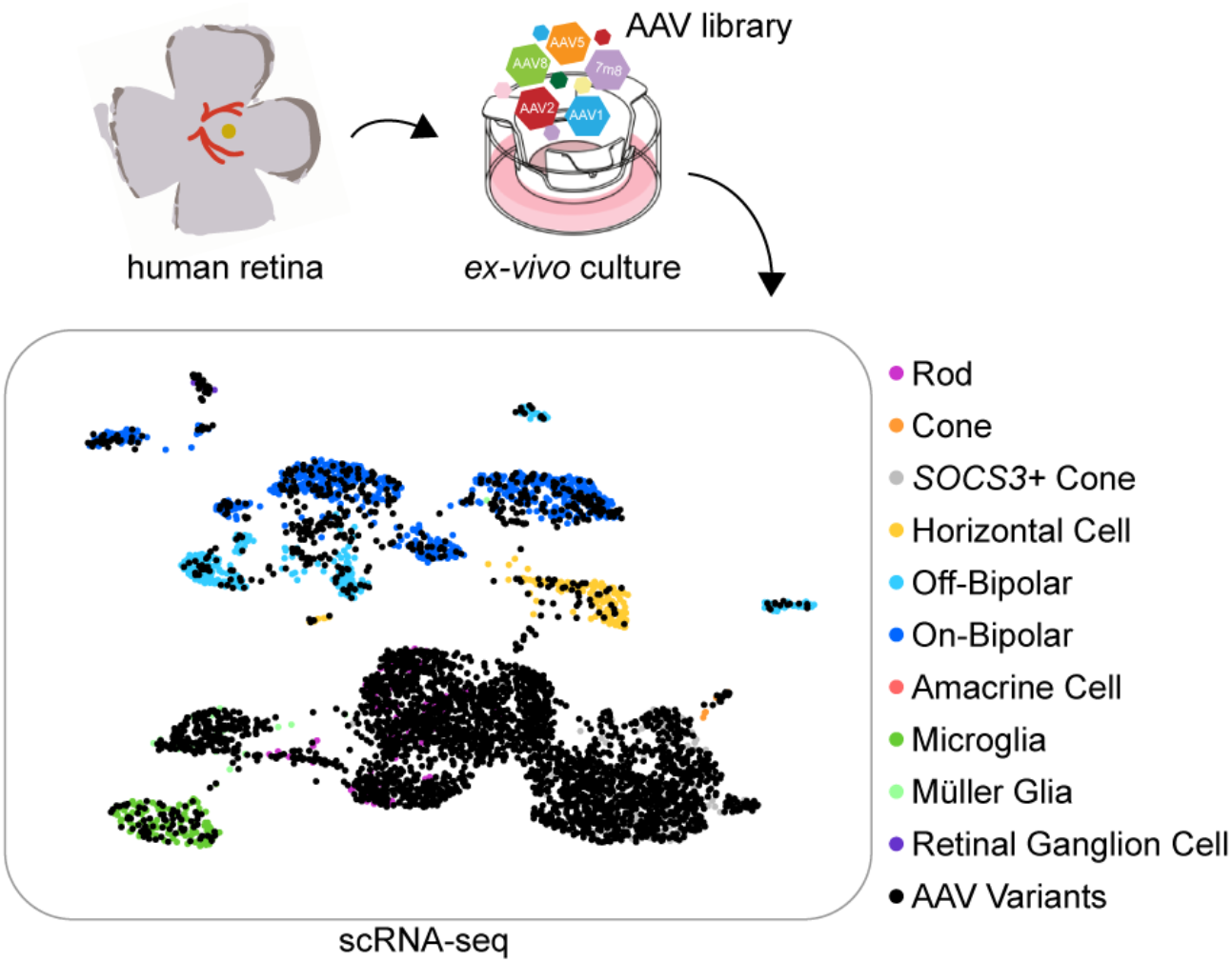

## Introduction

The FDA approval of adeno-associated virus (AAV) for treatment of Leber congenital amaurosis is a milestone in the field of gene therapy, ^1^ and a variety of AAV-mediated gene therapies for a wide range of diseases such as inherited retinal dystrophies, neuromuscular disorders, hemophilia and inherited metabolic disorders are at an advanced stage of clinical development.^2,3^ Efficient gene delivery is key to the success of each of these gene therapy approaches. Levels of transgene expression are determined by the properties of the viral capsid, the promoter driving transgene expression, the injection route, and the timepoint of delivery.^4^

A variety of studies have shown that changes can be made to the structure of the protein capsid shell of the virus, and that reengineering of the viral capsid can result in AAVs with improved abilities, enabling the application of lower doses of AAV, and diminished risks of adverse effects.^5^ Mutation of surface-exposed tyrosine, serine, and threonine residues resulted in viruses with increased infectivity, and protection of the AAV capsid from degradation. And, directed evolution (DE) approaches have resulted in the development of newly engineered AAVs such as 7m8, which has an improved ability to bypass structural barriers in the retina and infect photoreceptors.^6^ A number of clinical trials using these next-generation engineered viral vectors are ongoing (ClinicalTrials.gov; NCT03316560, NCT02416622, and NCT03748784).

It is essential to accurately quantify transgene expression levels and cell-type tropism of vectors to allow for the optimal selection of capsids for clinical translation. Mice have, to date, been the mostly widely used preclinical animal model for such studies. However, due to the anatomical and structural differences, AAV expression patterns differ dramatically in the retina of mouse and non-human primate (NHP), the large animal model with the highest similarity to humans.^6^ The quantification and validation of AAVs in NHPs, however, has, in the past, been difficult due to variability between animals, as well as the cost and ethical burdens associated with such work. Therefore, we recently established a single-cell RNA-seq AAV engineering (scAAVengr) pipeline for rapid evaluation of transgene expression, which allows direct, head-to-head comparison of multiple vectors across all cell types, in parallel, in the same animals.^7^ We first validated the scAAVengr pipeline in NHP retina.

Although the NHP retina is highly similar to human’s, a direct correlation in the distribution and specificity of AAVs to human retina is yet unproven. It’s therefore important to validate new AAV capsids in human tissue prior to clinical application. In this study, we applied the scAAVengr pipeline to human retinal explants, in order to quantify the tropism of 18 wildtype and recently engineered AAV capsids with single-cell resolution. Simultaneously, we compared the expression profile of human *ex vivo* cultured retina to fresh retina. Our study further advances the understanding of the retinal *ex vivo* culture model, and provides detailed information about the performance of leading clinically-relevant AAV serotypes in human retinal explants.

## Results

### Construction of the AAV library

In order to provide quantitative information about the tropism and efficiency of promising AAV variants in human retinal tissue, we screened an 18-member AAV library on human retinal explants using the scAAVengr pipeline (Figure 1). The AAV variants in the library included naturally occurring serotypes: AAV1, AAV2, AAV5, AAV8, AAV9, AAVrh10; tyrosine and threonine-mutated AAVs: AAV2-4YF, AAV2-4YFTV, AAV8-2YF, and AAV9-2YF;^8–10^ canine-derived DE variants K91, K912, K916, K94;^7^ NHP-derived DE variants NHP9, NHP26, and SCH/NHP26;^11^ and 7m8, a variant created through DE in mouse retina^6^ (Table 1). Each variant was packaged with a GFP transgene fused to a unique 25 bp barcode. The library was then created by pooling together the GFP-barcoded AAV variants, and the representation of each variant in the library was determined by deep sequencing (see methods).

**Figure 1.**
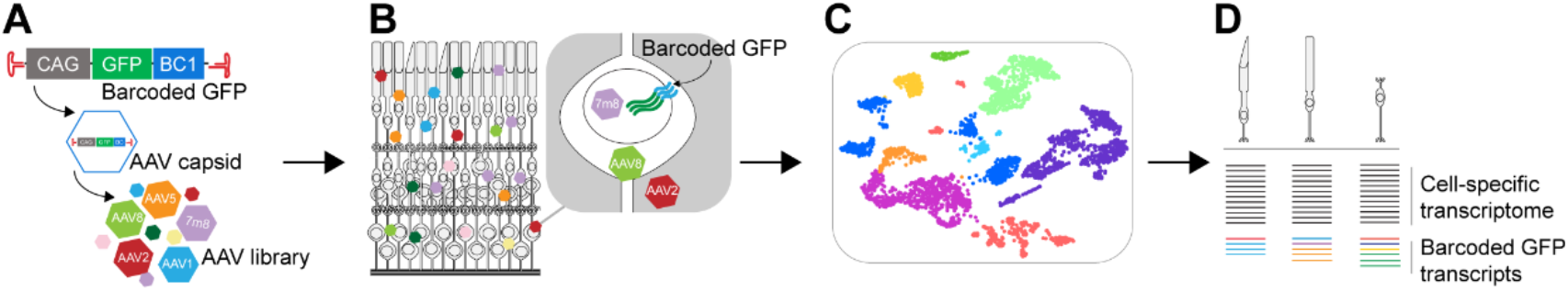
Illustration of the scAAVengr pipeline. (A) Generation of the AAV library. A CAG-GFP construct fused with a unique barcode was packaged into an AAV capsid. AAV variants containing unique barcodes were pooled and the representation of AAV’s in the library was quantified by deep sequencing. (B) AAV infection in the retina and transgene expression. Successfully infecting AAVs enter the nucleus and drive the expression of capsid-specific barcoded GFP mRNA in the infected cells. (C) The AAV-infected retina is processed for single-cell RNA-sequencing. Retinal tissue is dissociated and the transcriptomes of single cells are sequenced. UMAP plots are used to visualize clusters of retinal cells, and the cell types of clusters are identified based on the expression of cell-type specific marker genes. (D) The barcoded GFP transcripts are quantified, allowing for the tropism and infectivity of AAV variants to be evaluated across cell types with single-cell resolution.

**Table 1.**
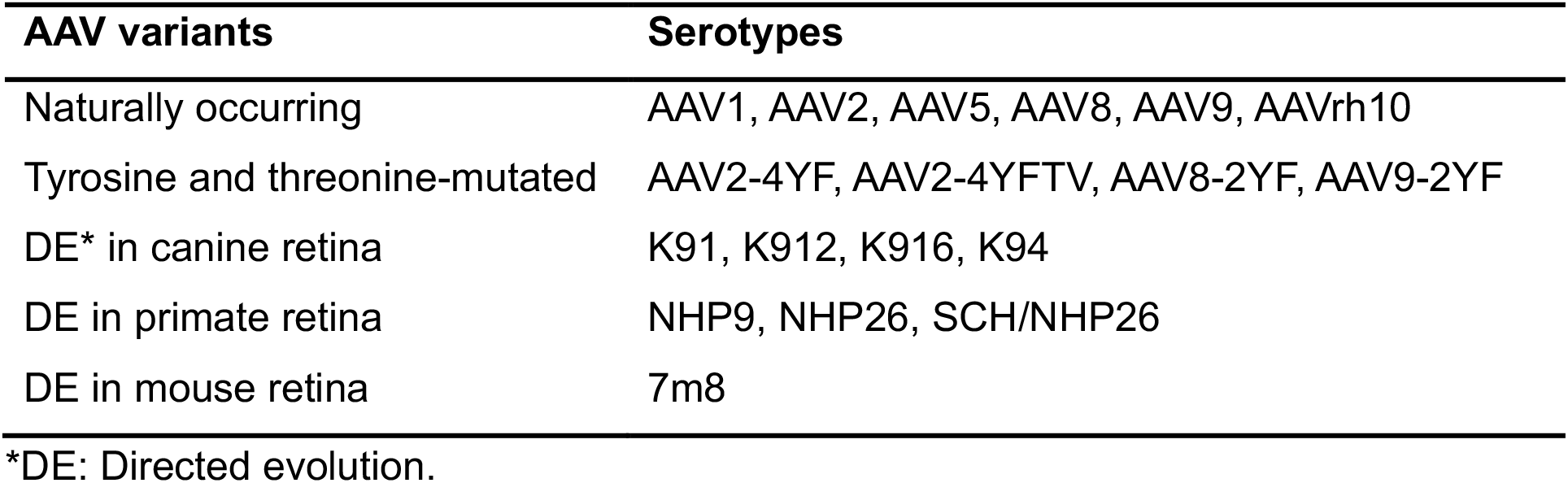

| AAV variants | Serotypes |
| --- | --- |
| Naturally occurring | AAV1, AAV2, AAV5, AAV8, AAV9, AAVrh10 |
| Tyrosine and threonine-mutated | AAV2-4YF, AAV2-4YFTV, AAV8-2YF, AAV9-2YF |
| DE* in canine retina | K91, K912, K916, K94 |
| DE in primate retina | NHP9, NHP26, SCH/NHP26 |
| DE in mouse retina | 7m8 |

\*DE: Directed evolution.

### Tropism of AAV variants in human retinal explants

The macular, mid-peripheral and peripheral regions from the temporal quadrant of human donor#1 retina were dissected and cultured *ex vivo* with the photoreceptor side facing down on a transmembrane placed in a culture well (Figure 2A-B). Retinal explants from the left eye and right eye were cultured in separate wells. AAV incubation was performed 1 day after culturing with 10μl of AAV library applied to the surface of each explant, and repeated every second day with the complete replacement of the medium. A total of 4.94E+11 vgs of AAV library in a volume of ~120ul was added to each culture well over the course of 8 days of AAV incubation.

**Figure 2.**
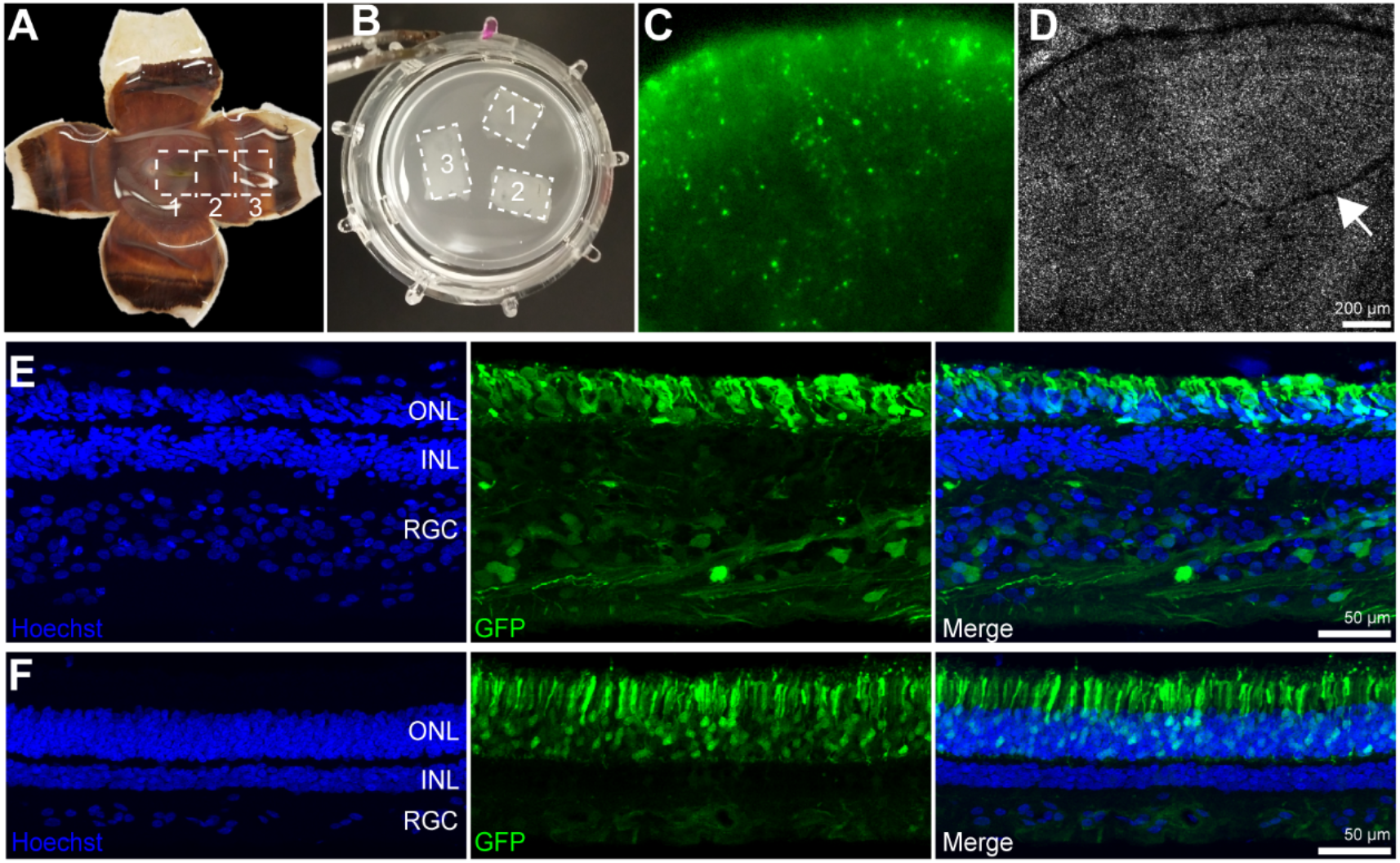
Human retinal explant culture and the tropism of the 18-member AAV library. (A) Dissection of the macular, mid-peripheral and peripheral retina from a postmortem donor eye. (B) Retinal explants were cultured on a transmembrane in a culture insert. (C) FITC channel and (D) trans channel. Images were captured at day 8 of AAV incubation with the explant still on the transmembrane. GFP expression was the strongest on the edge of the retinal explant, regardless of the presence of blood vessels (arrow). (E) Macular retinal explant. (F) Peripheral retinal explant. GFP expression was observed across the retinal layers and was most prominent in the outer retina.

GFP expression appeared on the edge of the retinal explant at day 2 of AAV incubation, with an additional area of increased expression towards the center of the retinal explant. This expression pattern was the same for retinal explants from all anatomical locations, including macular, mid-peripheral and peripheral retina, regardless of the presence of anatomical structures such as major blood vessels or the fovea (Figure 2C-D and Figure S1). On day 8 of AAV incubation (day 9 of the explant culture), the highest infected region (edges) of each explant was dissected and collected. Each explant (total 6 explants representing 3 regions from left and right eye combined) was processed individually for single-cell RNA seq (scRNA-seq). The single-cell suspension of macular, mid-peripheral and peripheral retinal explants from the same eye were then combined, and GFP positive cells were enriched by FACS (2 FACS-sorted samples representing all 3 regions from left and right eye) and processed for scRNA-seq. The remainder of the explant was fixed for imaging. Retinal explant cross sections showed that the basic structure of the retina was retained, with the presence of the retinal ganglion cell layer, inner nuclear and outer nuclear layers apparent in imaging. GFP was expressed across the retinal layers with the strongest expression in photoreceptors in the outer retina (Figure 2E-F).

### scRNA-seq quantification of AAV efficiency across cell types in human retinal explants

Single-cell cDNA libraries created from macular, mid-peripheral, peripheral retina regions, and FACS-sorted cells from all retinal regions combined, were deep sequenced and transcripts were aligned and quantified using STARsolo.^12^ Data from the same anatomical location in the left eye and right eye were combined, empty droplets were removed,^13^ doublets were removed,^14^ transcripts were normalized,^15^ and imputation was employed to improve the sparsity of single-cell data.^16^ Cells were clustered in a low dimensional space and cell type labels were identified for each cluster based on the most significant differentially expressed retinal cell type marker genes. The numbers of cells analyzed, after filtering, were: macula 4,860; mid-periphery: 2,480; periphery: 3,131; FACS: 3,162. Barcoded GFP transcripts originating from the various AAVs were then quantified using Salmon^17^ and mapped to the identified cell types using the associated single-cell barcodes (Figure 3, and refer to methods for details).

**Figure 3.**
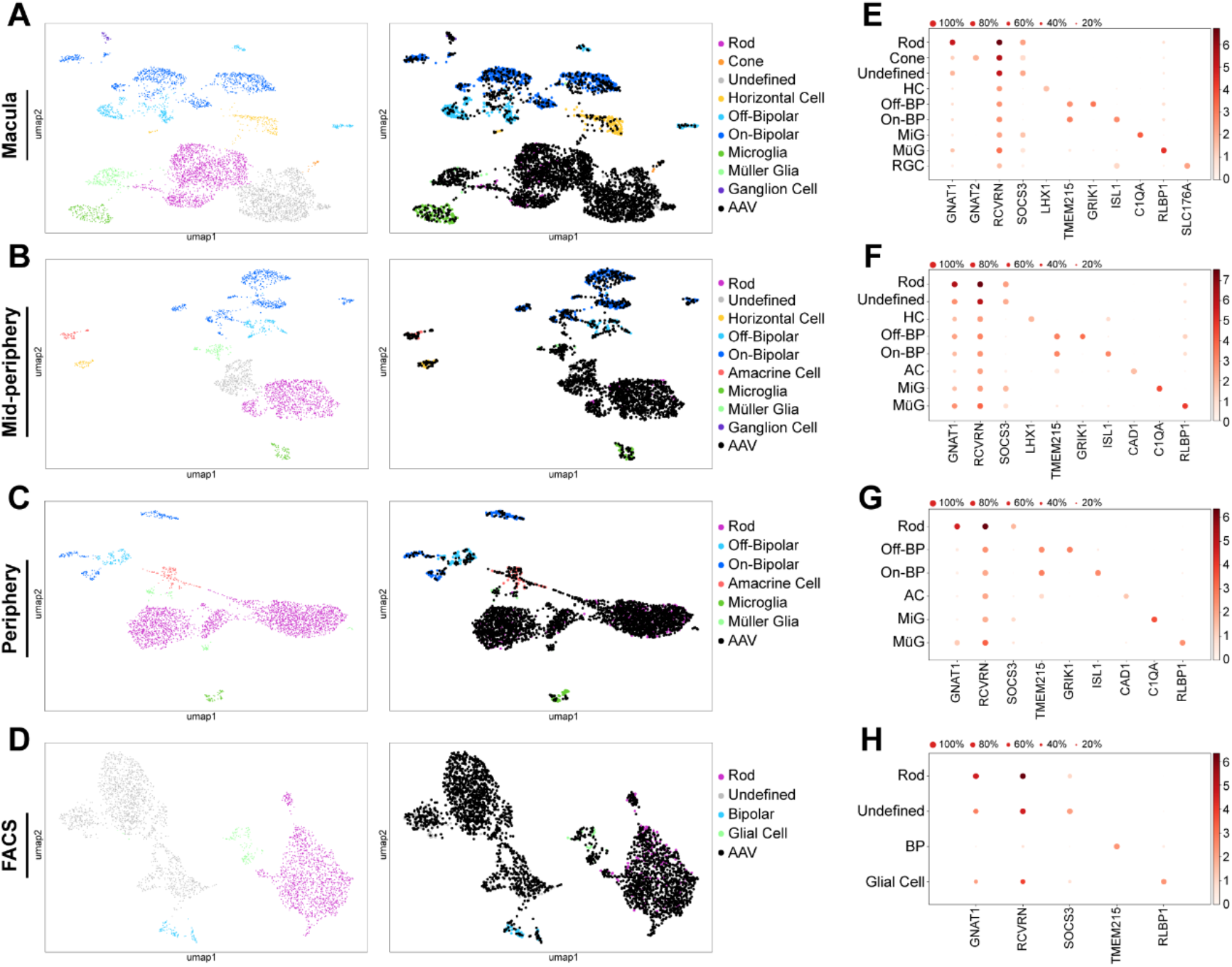
Cell clustering and identification of AAV infected cells. (A-D) UMAP plots of retinal cells, with and without AAV expression overlayed, from macular, mid-peripheral, peripheral and FACS samples. The color of the cell cluster indicates cell type, and AAV-infected cells are shown in black. (E-H) Heat maps of marker genes used for cell type identification. Each plot was generated with 2 samples from the left eye and right eye.

Clusters of major retinal cell types including rods, cones, bipolar cells, ganglion cells, amacrine cells, horizontal cells and glial cells were identified in the retinal explant samples (Figure 3). This is in agreement with scRNA-seq data performed with fresh retina from donor#2 (Figure S2A-C), and as previously reported.^18^ However, the cone population was only identified in the macular retinal explant samples, with the number of cones lower than expected. The number of cones identified did not correlate with histological imaging. Cones were labeled with Peanut Agglutinin (PNA), but were swollen and significantly shortened in the retinal explants compared to freshly fixed retina from the same eye (Figure 4 A-D). This was especially apparent at the edge of the explants, where tissue was collected for scRNA-seq (Figure 4 E-G). UMAP plots revealed a population of cells that did not express the cell-specific markers used for cell-type identification, but had high expression of GFP transcripts (Figure 3, undefined clusters). This cluster maps closely to the rod and cone population, and was enriched in FACS-sorted samples, which were largely populated with photoreceptors (Figure 3D). We thus hypothesized that this may be a group of cones with an altered transcriptome profile. In addition to *GNAT2*, we quantified expression of other cone-specific gene markers including *ARR3*, *GNGT2*, *OPN1LW*, *OPN1SW*, *OPN1MW*, *PDE6H*, none of which were significantly differentially expressed in the undefined cluster. Interestingly, as in rods and cones in the fresh retina samples, this undefined cluster had a high expression level of Recoverin (*RCVRN*), a pan-photoreceptor marker (Figure 3 and Figure S2). In addition, in this undefined cluster, we found high expression of suppressor of cytokine signaling 3 (*SOCS3*), a well-known negative regulator of cytokine signaling which is involved in many cellular processes, including inflammation and cell death. In fresh retina samples, in contrast, the expression of *SOCS3* is limited to the microglia (Figure S2). *SOCS3* has been considered as an indicator of retinal stress and shown to be upregulated during retinal degeneration.^19^ This evidence strongly suggests that the undefined cell cluster represents degenerating cones lacking cone-specific markers. While cones were only identified in the macular samples, this *SOCS3*+ cone population was the largest in the macular samples, reduced in size in the mid-peripheral retina samples, and absent from the peripheral retina samples. This may result from the relative cone densities, which vary according to eccentricity and are most concentrated in the macular retina, and decrease in density towards the periphery of the retina.

**Figure 4.**
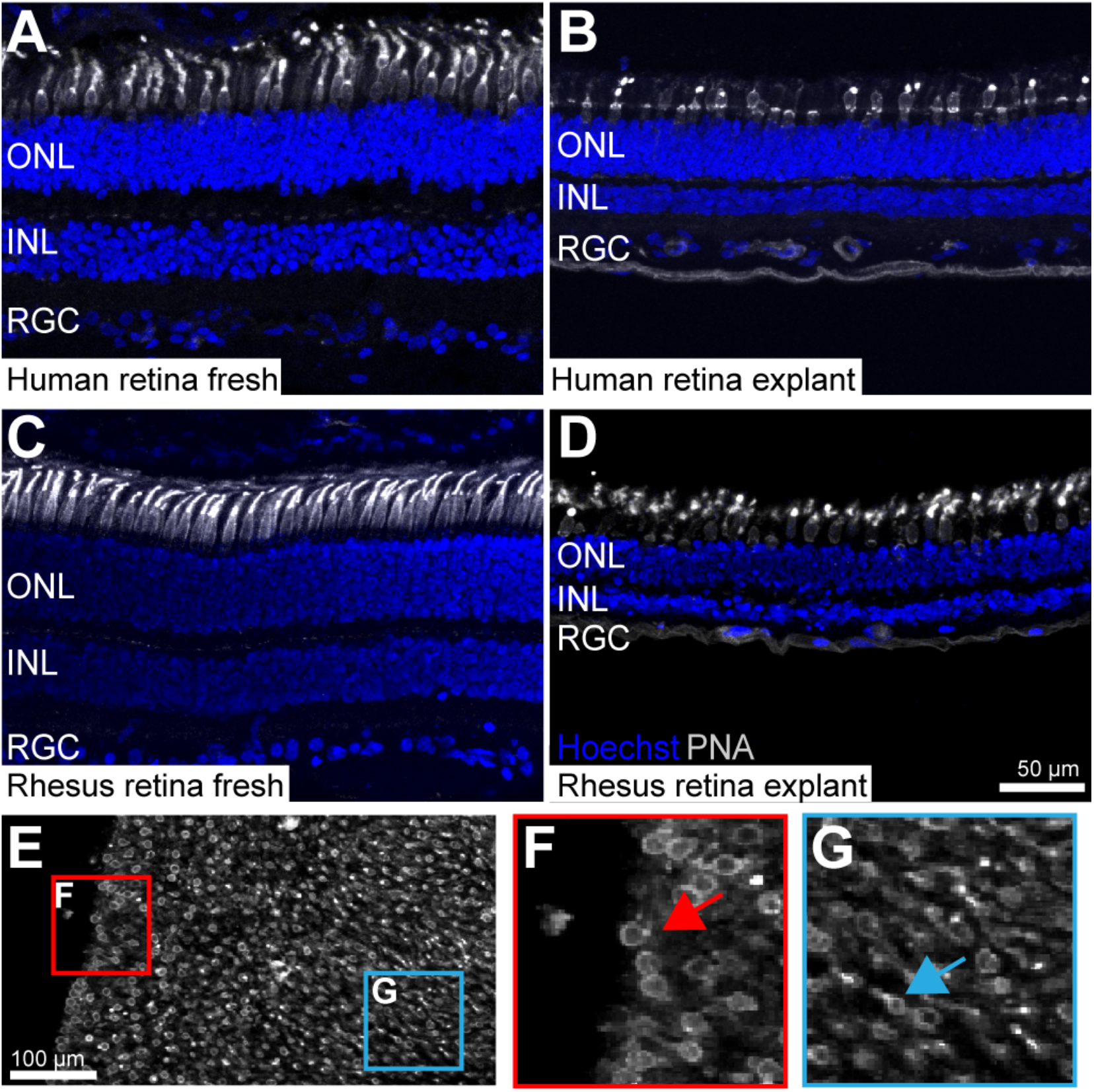
Unhealthy cones in retinal explants. (A) Freshly fixed human retina. (B) Human retinal explant at day 9 of culturing. (C) Freshly fixed rhesus retina. (D) Rhesus retinal explant at day 9 of culturing. (E-G) A flatmounted human retinal explant at day 9 of culturing. Unhealthier cones (rounded, swollen) are observed at the edge, while the morphology of cones is better preserved (elongated) at the center.

The performance of individual AAV serotypes was then evaluated across cell types using three metrics: the absolute number of cells infected by each serotype (Figure 5A); the percent of each cell type infected by each serotype (Figure 5B); and the level of transgene expression mediated by each serotype in the infected cells (Figure 5C). Each of these metrics was corrected by the AAV dilution factors determined by deep sequencing of the library. Heat maps of these metrics revealed that AAV variants engineered through DE outperformed naturally occurring AAVs as well as tyrosine and threonine-mutated AAVs across cell types and in explants from all three anatomical locations. Photoreceptors were the most efficiently infected cell type in all three regions, with the highest number of cells infected, the highest percent of cells infected, and the highest mean transcripts of infected cells. This is in agreement with GFP expression observed in retinal cross sections (Figure 2E-F). Also, the majority of cells identified in FACS-sorted samples, where the GFP positive cells were enriched, were rods and *SOCS3*+ cones (Figure 3D).

**Figure 5.**
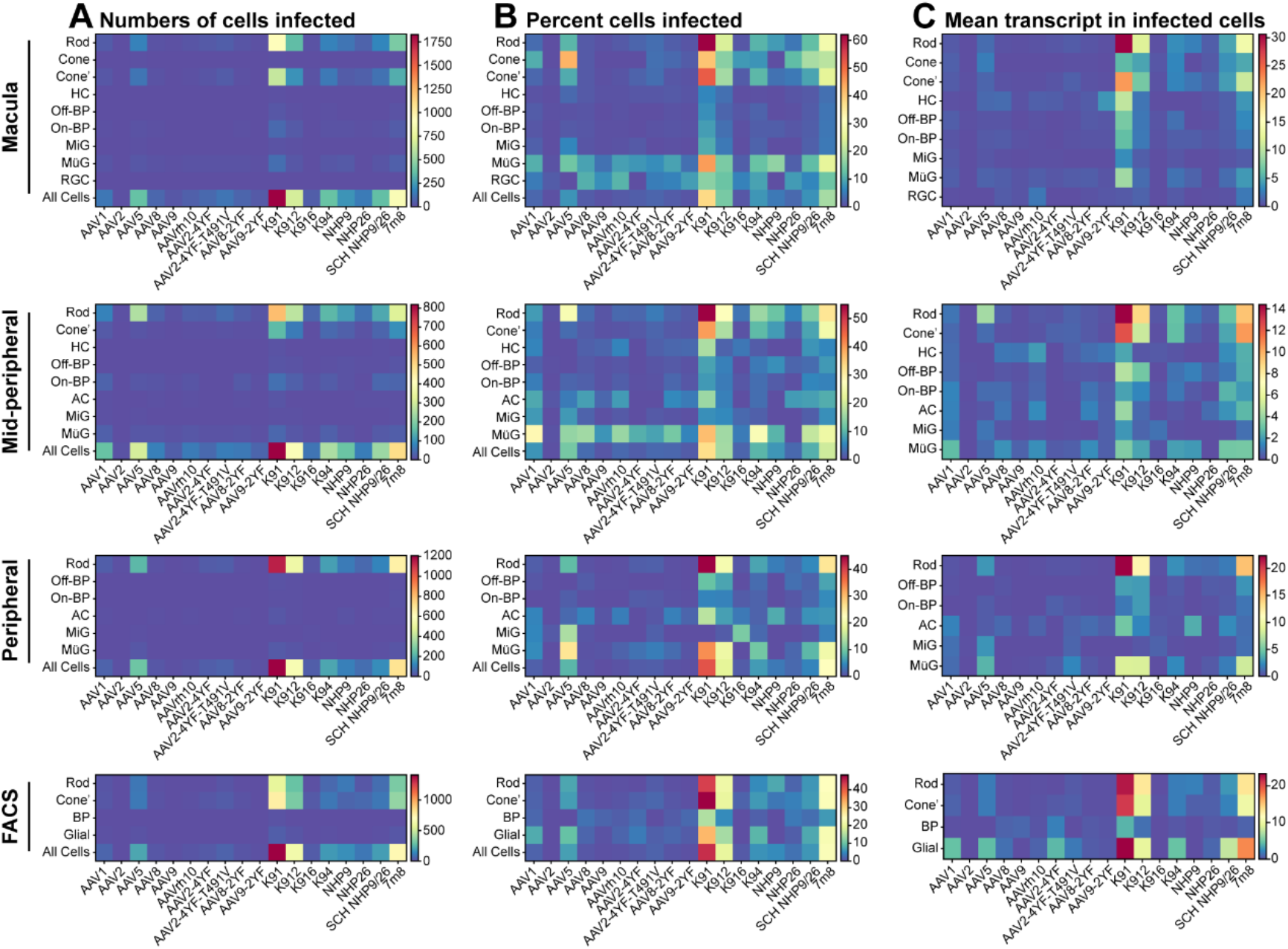
Quantitative comparison of AAV variant expression across retinal cell types. (A) Numbers of cells infected by each AAV variant. (B) Percent of each cell type infected by each AAV variant. (C) Level of transgene expression driven by each serotype in the infected cells. Data are shown as mean transcripts per cell/100,000 transcripts. All data are corrected by the AAV dilution factor. Each plot was generated with 2 samples from the left eye and right eye. Cone’ = *SOCS3*+ cones. HC = Horizontal Cell; BP = Bipolar Cell; AC = Amacrine Cell; MiG = Microglia; MüG = Müller Glia; RGC = Retinal Ganglion Cell.

Next, in order to rank the best performing pan-retinal serotypes by cell type, variants were plotted by the mean transcripts per cell in infected cells vs. the percent cells infected for each AAV serotype (Figure 6). Across retinal cell types and regions, of the canine-derived variants, K91 outperformed other engineered serotypes. Of the primate-derived variants, SCH NHP9/26 outperformed other variants. Among all the AAVs, canine-derived variant K91 showed the highest infectivity at all retinal regions, closely followed by canine-derived variants K912 and mouse-derived variant 7m8. While the naturally occurring serotypes are overall less efficient, AAV5 was the most efficient of the naturally occurring serotypes tested in photoreceptors (Figure 5 and 6).

**Figure 6.**
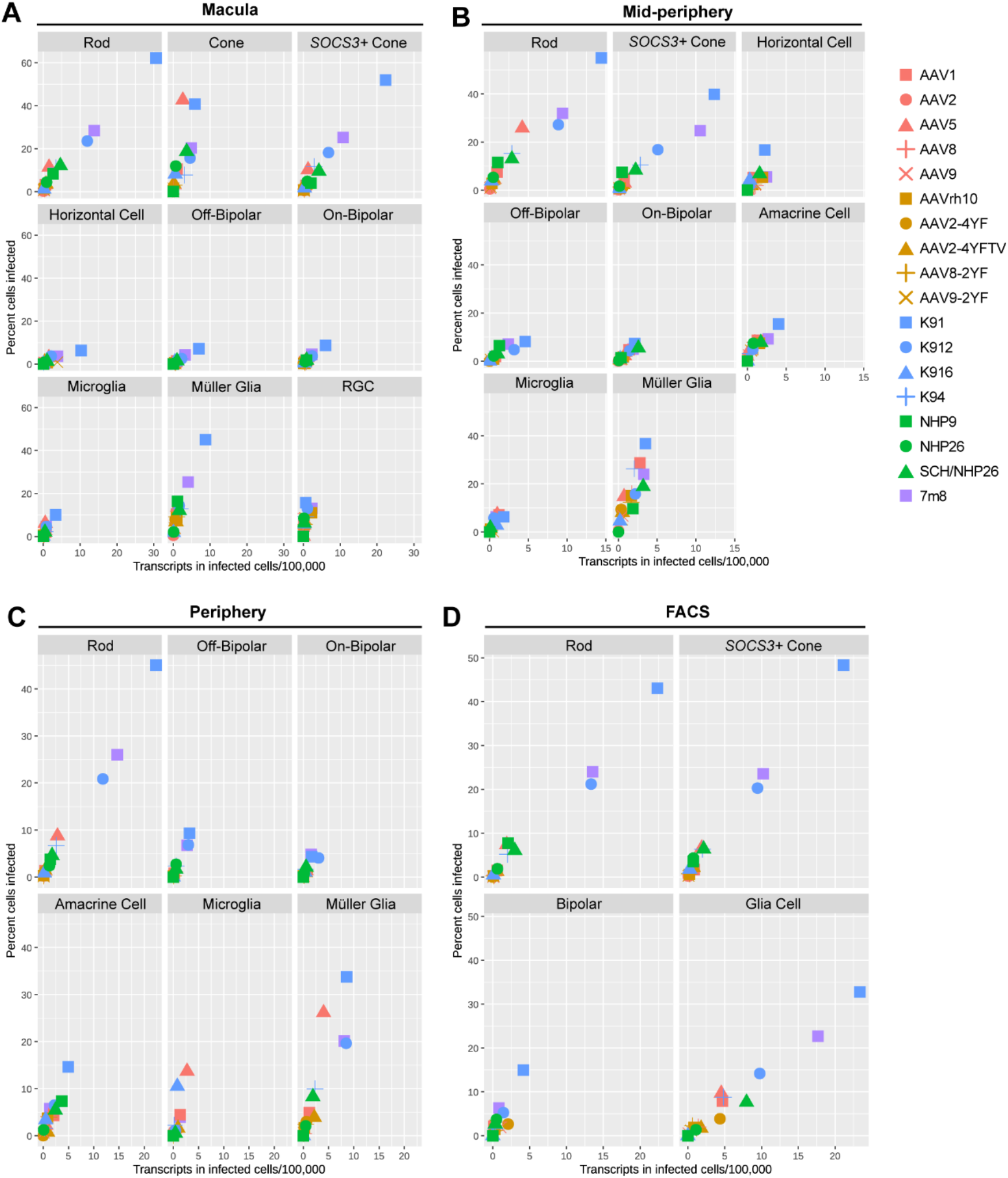
Serotype performance across retinal regions. Scatter plots show the number of transcripts in infected cells per 100,000 transcripts vs. the percent of cells infected for each serotype. Best performing variants that infect the highest number of cells and express the most GFP transcripts appear towards the upper right corner of each plot. All data are corrected by the AAV dilution factor. Each plot was generated with 2 samples from the left eye and right eye. RGC = Retinal Ganglion Cell.

We have previously shown that multiple AAVs can infect a single cell.^7^ In order to understand the dynamics of AAV infection in human retinal explants, upset plots with the number of cells infected by a particular combination of AAVs (the intersection size) and the number of cells infected by a particular serotype (the set size) were generated (Figure 7). As many as 14 serotypes simultaneously infected a single cell. As revealed by the intersection size, more cells were infected by a single high-performing serotype (K91, K912, and 7m8. Figure 7, black dots section) or by the combinations of two (Figure 7, red lines section) or three (Figure 7, yellow lines section) serotypes. Base on the set size, without AAV dilution factor correction, K912 infected the greatest number of cells, followed by K91 and 7m8.

**Figure 7.**
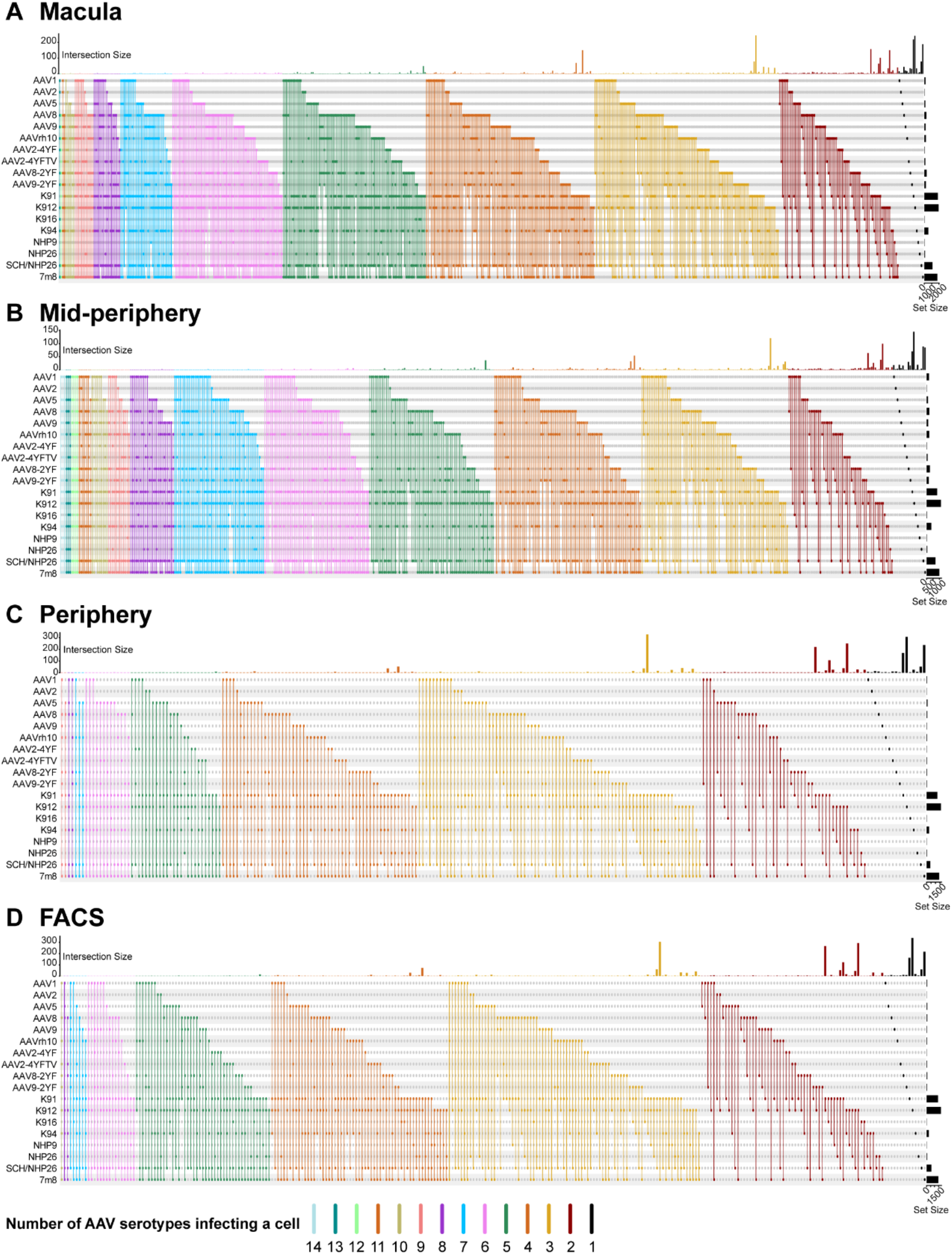
Cells infected by multiple AAV serotypes. The number of cells infected by a particular combination of AAVs (the intersection size) is shown in bar graphs across the top of each plot, and the number of cells infected by a particular serotype (the set size) is shown across the right-hand Y-axis. Dots and connecting vertical lines indicate the serotype and number of variants infecting a single cell. Lines are colored according to the number of AAV variants in the subset.

### Validation of top-performing AAV serotypes

In order to validate the top-performing AAV serotypes, and to compare AAV performance *ex vivo* and *in vivo*, we tested individual AAV variants in NHP and human retinal explants. CAG-GFP constructs were packaged into K91 and K912. K91 was the best performing serotype in human retinal explants, but this variant has previously been shown to underperform compared to other variants *in vivo* via intravitreal injection in NHP retina.^7^ K912 was the second best performing serotype in human retinal explants, and the top-performing serotype among the set tested *in vivo* in NHP retina.^7^

Additionally, to better understand whether the AAV tropism is affected by the orientation of the explants in the culture dish (the medium containing AAVs contacts the side of explants attached to transmembrane but does not submerge the explants), one set of retinal explants was cultured with the photoreceptor side attached to the transmembrane (PR-down), and another set of explants was cultured with the RGC side attached to the transmembrane and photoreceptor side facing upwards (PR-up). The PR-up retinas were infected with either K91-CAG-GFP or K912-CAG-GFP, and the PR-down explants were infected with K912-CAG-GFP. K91-CAG-GFP and K912-CAG-GFP were titer matched, and a total of 2.45E+10 vgs of AAV in a volume of 120 μl were applied to each group as previously described during 8 days of AAV incubation (each group consists of 3 explants, which were collected from central, mid-peripheral and peripheral retina).

On day 2 of AAV infection, GFP expression was first observed on the edge of explants in the K91 PR-down, K912 PR-down and K912 PR-up groups. The GFP expression increased in intensity and area with time and plateaued in intensity by day 8 of AAV infection (Figure 8A-C). The retinal explants were then fixed and sectioned. In the PR-down groups, both K91 and K912 drove efficient GFP expression across retinal layers, with the highest expression observed in the outer retina, in accordance with data from scAAVengr analysis in the human retinal explants (Figure 8D-E). Interestingly, the K912 infected PR-up explants also showed GFP expression mainly in the outer retina (Figure 8 F). The tropism of K912 in rhesus retinal explant was similar in both culture orientations, suggesting that the orientation of the explant in culture does not influence tropism, and that K912 has high affinity for photoreceptors.

K912, which outperformed other variants both *in vivo* in NHP retina and *ex vivo* in human retinal explants, was further validated in retinal explants from another human donor (donor#3). 7m8, which infects human explants efficiently ^20–22^ and has been used in clinical trials,^23^ and naturally occurring AAV2, the first clinically approved serotype, were tested in parallel. The *ex vivo* culture of central, mid-peripheral and peripheral retina and AAV application were performed as previously described. A total volume of 120μl of K912-CAG-GFP (1.26E+10 vgs), 7m8-CAG-GFP (5.58E+11 vgs), and AAV2-CAG-GFP (1.33E+10 vgs) were applied during 8 days of AAV incubation. K912 and 7m8-infected explants started to show GFP expression on the edge of the explants at day 2 while AAV2 (in a similar dose as K912)-infected retina showed GFP expression at day 4 (Figure7 G-I). In retinal cross sections, the strongest GFP expression was observed in the outer retina for all three serotypes. K912 and 7m8 drove significantly stronger GFP expression than AAV2 (Figure 8 J-L).

**Figure 8.**
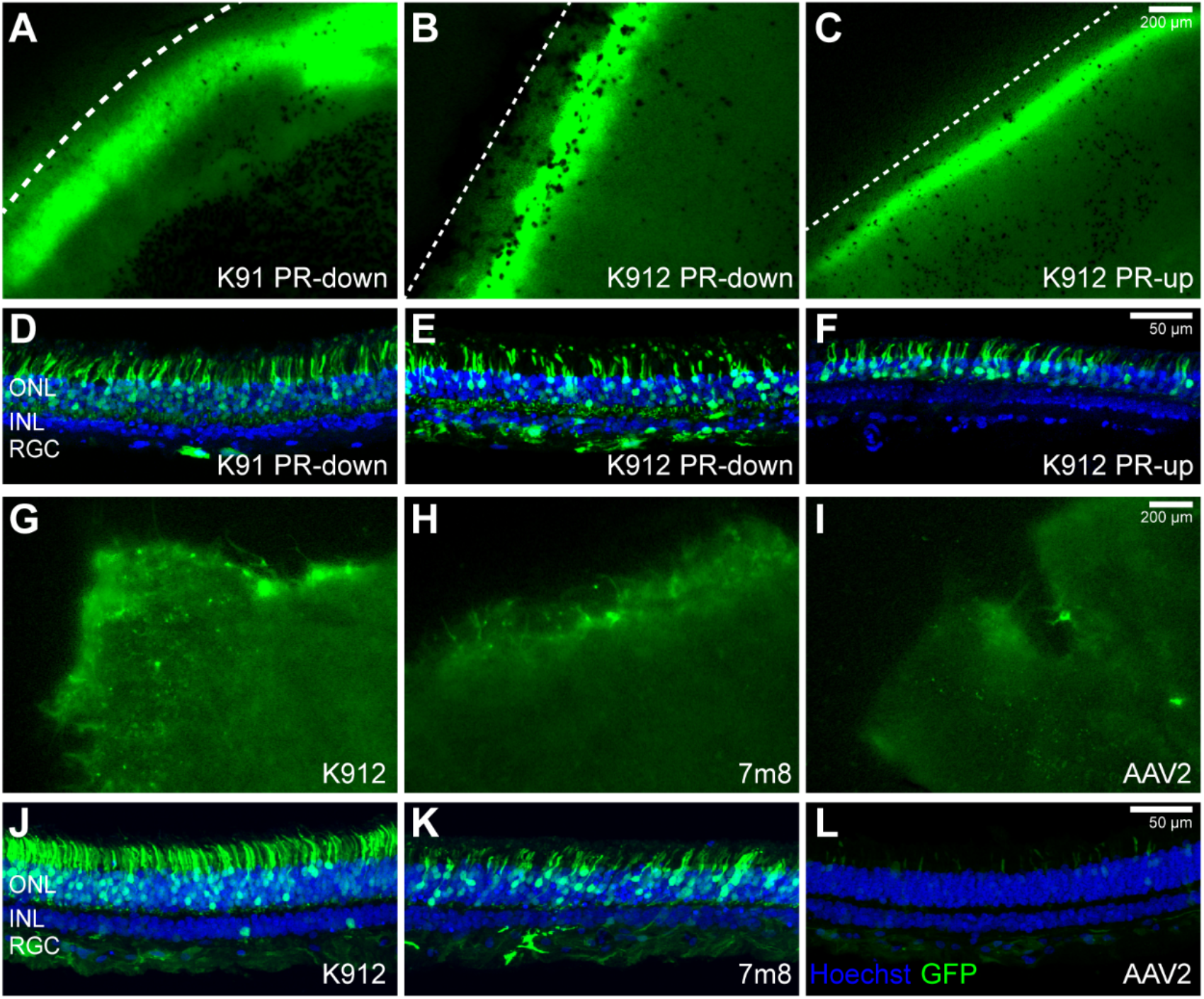
Validation of individual AAV serotypes. (A-C) GFP expression of K91 and K912-infected rhesus retinal explants 8 days post infection. Dotted line, edge of the retinal explants. (D-F) GFP expression in cross sections from the K91 and K912-infected rhesus retinal explants 8 days post infection. (G-I) Onset of GFP expression in the K912, 7m8, and AAV2-infected human retinal explants. (J-L) GFP expression in the K912, 7m8, and AAV2-infected human retinal explant cross sections 8 days post infection. (J-L) were imaged with the same acquisition parameters. Each group has 3 explants from central, mid-peripheral and peripheral retina.

## Discussion

*Ex vivo* retinal culture is the only available transduction model of mature human retinal tissue, and it has been widely used in retinal research, bridging the gap between *in vitro* cell/organoid culture and *in vivo* experimental animals. In this study, we applied the scAAVengr pipeline to human retinal explants, allowing for head-to-head evaluation of the tropism of 18 AAV capsid variants with single-cell resolution. These experiments provide quantitative information about the leading serotypes currently in development for clinical application. Overall, we found that AAV capsids engineered through DE have greater infectivity than naturally occurring AAV variants or tyrosine and threonine-mutated AAVs in human retinal explants, across cell types and in all three anatomical locations, which is in accordance with previous observation *in vivo* in NHPs.^7^ K91 was the best performing serotype in human retinal explants, closely followed by K912 and 7m8. All three of these serotypes mediated efficient gene expression in human and rhesus retinal explants, which was confirmed through validation of individual variants.

In a previous scAAVengr experiment, performed using intravitreal injections in NHP retina *in vivo*, K912 was identified as a top performing panretinal virus, but K91 did not outperform parental serotypes.^7^ In our *ex vivo* human retinal culture, in contrast, K91 had better performance than K912 as quantified by the scAAVengr pipeline. To rule out the possibility that the different performance of K91 in human *ex vivo* tissue is due to cross-species differences, K91-CAG-GFP was packaged separately and used to infect NHP retinal explants. Transfection with K91-CAG-GFP drove high level of transgene expression in NHP retinal explants, indicating that the improved performance of K91 is a result of the *ex vivo* culture system, rather than any species specificity.

The lower efficiency of K91 *in vivo* via intravitreal injection may be due to lower ability to bypass the anatomical barriers that exist *in vivo* but are not intact in the *ex vivo* culture system. Retinal tissue in *ex vivo* culture maintains its normal gross morphological structure, as well as heterogeneous cell populations.^24,25^ However, anatomical structures including the vitreous and inner limiting membrane, barriers that restrict AAV diffusion *in vivo* are not intact, especially on the edge of the explants.^26^ In explants from all anatomical locations including macular, mid-peripheral and peripheral retina, AAVs showed higher expression levels near the edge of the explants than in the center of the explants, likely due to this lack of anatomical barriers at their edges allowing for easier accessibility of AAVs.

In our experiments, both the inner and outer retina of the explants have direct access to AAVs. The AAV suspension was added onto the RGC side of the explants, and then quickly diffused into the medium, which is in contact with the photoreceptor side of the explants. We therefore compared the expression pattern of AAVs in retinal explants with expression patterns from intravitreal and subretinal injections. *In vivo*, following intravitreal injection, the highest transgene expression is observed in the foveola and in a perifoveal ring of retinal ganglion cells, and in the peripheral retina in punctate areas near blood vessels.^6^ In retinal explants, no regional differences of expression patterns were observed, and we did not observe higher expression of GFP in the foveal pit or near blood vessels. This lack of AAV transfection in the foveola and around blood vessels may also be due to differences in the accessibility or presence of AAV receptors in these regions, and further histological studies will be required to better understand this differing expression pattern.

Subretinal injections *in vivo* result in strong transgene expression in photoreceptors under the injection bleb. Several AAV serotypes mediate high levels of transgene expression in photoreceptors following subretinal injection *in vivo.* AAV5, for example, although it has low efficiency via intravitreal injection, drives fast-onset and efficient expression in photoreceptors when administered into the subretinal space.^4^ As revealed by histological analysis, and with the scAAVengr workflow, the strongest GFP expression in retinal explants, driven by a ubiquitous CAG promoter, was observed in photoreceptors, with both serotypes in the AAV library, including AAV5, and the individually tested serotypes. The same expression pattern in retinal explants has previously been reported by other groups.^27,28^

To better understand if the high AAV infectivity in photoreceptors is a result of easier accessibility due to the orientation of the *ex vivo* culture system, we cultured rhesus retinal explants with either photoreceptors facing down to the transmembrane or facing up, and infected with AAV K912-CAG-GFP, a high-performing serotype in both *ex vivo* human retina and *in vivo* NHP retina.^7^ The highest expression levels of GFP were observed in the outer retina in both culture orientations. This indicates that the tropism of AAVs in retinal explant culture is more comparable to subretinal injections than intravitreal injections *in vivo* regardless of culture orientation.^29^ Therefore, while retinal explants may provide a valuable system in which to confirm AAV infectivity for human retinal cells, the usefulness of *ex vivo* culture may be limited for the prediction of tropism via intravitreal injection *in vivo*. Additionally, both the scAAVengr pipeline and individual validation showed that K91 has high affinity for photoreceptors in human and NHP retinal explants, and it may be of interest to further evaluate the potential of K91 for subretinal delivery *in vivo*.

Although human retinal explant culture is an important preclinical model, a significant drawback of this model is the limited survival window.^30^ As seen through histological analysis, all retinal cell types degenerate over time.^24^ However, it is not well understood how retinal *ex vivo* culture affects the expression profiles of retinal cells compared to fresh retina. Here, for the first time, we evaluated the condition of the human retinal explant culture at the single-cell level. Under the conditions tested here, 9 days after culturing, the major retinal cell types expressed similar retinal cell type marker genes as the fresh retina, except for the cone population, which showed an altered expression profile with the loss of cone-specific genes and upregulation of *SOCS3*, an indicator of retinal stress. While *SOCS3* is only expressed in microglia in fresh retina, upregulation of this gene was observed in rods and cones in the retinal explant. Previous studies have also noted that photoreceptors are the fastest deteriorating cell types in *ex vivo* culture at the histological level, and that the degeneration pattern is comparable to retinal injury and degenerative diseases.^24,31^ The fact that AAVs efficiently infect degenerating photoreceptors in human retinal explants suggests that they may be promising for the treatment of retinal degeneration in patients.

In conclusion, we quantified the performance of 18 AAV serotypes in human retina *ex vivo* culture, and determined that K91, K912 and 7m8 are the most efficient serotypes. Traditionally, the efficiency of AAVs has been evaluated by the expression level of transgenes such as GFP via histological analysis, a method which lacks quantitative accuracy. Using the scAAVengr workflow, we were able to simultaneously compare the efficiency of multiple AAVs in the same retinal explants and in the same cells. These results provide detailed information about leading AAV vectors for the pre-clinical development of gene therapy approaches. This study also provides valuable insight to the benefits and drawbacks of using retinal explant culture for the evaluation of AAV serotypes. *Ex vivo* culture is a useful model for confirming the infectivity of AAV in human retina, yet it may not be the best system in which to determine the tropism of AAV serotypes *in vivo*, due to the discrepancies in infection patterns between *in vivo* and *ex vivo* models.

## Methods

### AAV production and quantification

A set of constructs containing unique 25 bp DNA barcodes after the stop codon of eGFP with a self-complementary CAG promoter, and flanked by ITRs, was packaged into individual AAV capsids. Each AAV serotype was packaged separately using a triple transfection method^32^using 293AAV cells (Cell Biolabs). AAVs were purified by iodixanol gradient ultracentrifugation, buffer exchanged and concentrated with Amicon Ultra-15 Centrifugal Filter Units (#UFC8100) in DPBS. Each variant was pooled and the titer of the virus was determined by quantitative PCR relative to a standard curve using ITR-binding primers or by using a QuickTiter AAV Quantitation Kit (Cell Biolabs). The relative titer of each variant in the pooled AAV library was confirmed by Illumina MiSeq sequencing (primer sequences are provided in Table S1).

### Postmortem human eyes

All experiments were performed with approval and oversight from the University of Pittsburgh Committee for Oversight of Research and Clinical Training Involving Decedents. Eyes from postmortem donors was obtained through the Center for Organ Recovery & Education. Donor#1 20-year-old male, donor#2 58-year-old female, donor#3 45-year-old male, no history of retinal disease.

### Rhesus Macaque

A 3-year-old male rhesus macaque was housed under standard 12-h light/12-h dark conditions. For euthanasia, the animal was initially sedated with ketamine (15 mg/kg IM), and then perfused through the circulatory system with 3-4 liters of ice cold artificial cerebrospinal fluid (ACSF; 124 mM NaCl, 5 mM KCl, 2 mM MgSO4, 2 mM CaCl2, 23 mM NaHCO3, 3 mM NaH2PO4, 10 mM glucose; pH 7.4, osmolarity 290–300 mOsm). Eye were then enucleated. All procedures were performed in accordance with the Association for Research in Vision and Ophthalmology statement for the Use of Animals in Ophthalmic and Vision Research. All animal experiments were approved by the University of Pittsburgh Institutional Animal Care and Use Committee (IACUC).

### Retinal explant culture

Eyes from postmortem donors/ rhesus were dissected within 30 minutes after surgical removal. The anterior segment was removed and eye cups were flat mounted. Around 5mm wide macular (central), mid-peripheral and peripheral retina was dissected and retinal tissue was carefully separated from RPE/choroid/sclera and transferred to cell culture inserts with 0.40 μm pore size (Thermo Fisher Scientific, #140640), with the photoreceptor side attached to the membrane (except for the PR-up group). Culture wells contained 3 retinal explants; macular (central), mid-peripheral and peripheral retina. The retinal explants were cultured in Neurobasal Plus Medium (Gibco, A35829-01) supplemented with 2% B-27 (Gibco, A35828-01), 5 μg/ml Plasmocin prophylactic (InvivoGen, #ant-mpp) and Penicillin-streptomycin (Genesee scientific, PSL01-100ML) at 37 °C, 5% CO_2_. In order to keep the retinal explant attached to the transmembrane, a volume of 1.3 mL of medium was added into each culture well, which is just enough for wetting the transmembrane but not enough to submerge the explant. Fresh medium was replaced at the second day of incubation and 10μl of AAV capsid library was applied dropwise onto the surface of each explant (30μl per culture well). Every second day, medium was completely replaced and AAV application was performed. A total of 120 μl of AAV suspension was applied per culture well during 8 days of AAV incubation. GFP expression in the retina was monitored using an ECHO Revolve fluorescence microscope. On the 8th day of AAV incubation, the 1mm edges of each explant, which had the highest level of GFP expression, were collected and underwent single-cell dissociation, FACS, and scRNA-seq. Retinal explants used for histological analysis were fixed with 4% paraformaldehyde (PFA) for 2-4 h.

### Single-cell dissociation of retina tissue

The 1mm edge of each retinal explant was placed in Hibernate solution (BrainBits, HE500) after dissection, and was then dissociated with an enzymatic and mechanical method using a MACS Neural Tissue Dissociation Kit for postnatal neurons (Miltenyi Biotec, 130-094-802) according to manufacturer’s recommendations. The macular, mid-peripheral and peripheral region from the temporal quadrant retina of human donor#2 was dissected, and the same anatomical regions from left and right eye were combined and immediately dissociated using the same method. The retinal single cells were resuspended in D-PBS with 0.1% BSA and processed immediately for scRNA-seq or FACS.

### Fluorescence-activated cell sorting (FACS)

The GFP-positive cells in the retinal explant cell suspensions were enriched using a MACS Tyto sorter (Miltenyi Biotec). GFP-positive cells were resuspended in D-PBS with 0.1% BSA and processed immediately for scRNA-seq.

### Single-cell RNA-Sequencing (scRNA-seq) and targeted gene enrichment

Following the manufacture’s protocol, using the Chromium Single Cell 3’ v3 kit (10x Genomics), single retinal cells were partitioned into Gel Beads in Emulsion (GEMs), mRNA was reverse transcribed to cDNA and barcoded with unique molecular identifiers (UMIs). The GEM emulsions were then broken and the cDNA was purified using DynaBeads. After PCR amplification, the cDNA was cleaned and further purified with SPRIselect reagent (Beckman Coulter, B23318). The final indexed library was constructed following fragmentation, end repair, A-tailing, adaptor ligation, and sample index PCR steps. The libraries were sequenced on Illumina S1 flow cells at the UPMC Genome Center. GFP transcripts with unique AAV barcodes were further enriched using a custom Targeted Gene Expression kit (10x Genomics) and the resulting library was sequenced with an Illumina MiSeq reagent nano kit v2 (300 cycles).

### scRNA-seq data processing and cell type identification

scRNA-seq data was aligned and cells were demultiplexed using STARsolo^12^ (v2.7). The GRCh38 reference genome (GCA_000001405.15) and its associated annotation file (NCBI RefSeq) were downloaded from UCSC and used for alignment. Empty droplets were removed using DropletUtils^13^ (v1.4.3, lower.prop=0.05). Doublets were removed using SCDS^14^ (v1.0.0), with a hybrid score cutoff of 1.3. Gene expression was normalized using Scran^15^ (v1.12.1) and imputation was achieved using ALRA^16^ (v1.0) to improve the sparsity of the scRNA-seq datasets.

The analysis of scRNA-seq datasets and cell type identification was performed with Scanpy^33^ (v1.4.4.post1). Specifically, the top 50 principal components of the gene expression matrix were calculated and used to compute the Euclidean distance between cells. Cells were embedded into a neighborhood graph for visualization using the UMAP algorithm, and Leiden clustering was performed on this graph to identify cell populations. Cell types were identified using a hypergeometric test using a list of known retinal cell type marker genes compiled from multiple sources ^34,35,36^. P-values were corrected for multiple hypothesis testing using the Bonferroni method and the cell type for each cluster was chosen based on the most significant marker gene intersection p-value (< 0.05). Clusters that did not meet the significance threshold for cell type identification were analyzed and annotated manually using the known gene markers.

### Quantification of AAV barcodes

AAV variants were identified using a unique 25 bp barcode at the end of the GFP gene for each variant. GFP was identified from these samples using two datasets: (1) whole transcriptome scRNA-seq data and (2) 10x targeted enrichment against GFP and other marker genes. Salmon^17^ (v0.9.1) was used for GFP transcript quantification and in-house scripts were used to identify the AAV barcode and map these reads back to their respective cells as previously described.^7^

### Immunohistochemistry

The retinal explants remained on the transmembrane and were fixed in 4% paraformaldehyde on ice for 2-4 h and dehydrated sequentially in 5%, 10%, 20%, and 40% sucrose solutions in PBS at room temperature (RT) (at least 30 min for each concentration). The transmembrane was cut around the retinal explant, and the retinal explants with the transmembrane were embedded in a 1:1 mixture of 40% sucrose in PBS: OCT compound (Fisher Scientific) in liquid nitrogen for cryosection. Sections of 14 μm were collected on glass slides. Cryosections were rehydrated with PBS for 10 min at RT followed by PBST (PBS plus 0.1% TritonX-100) for 30 min (retinal explant flatmounts were incubated in PBST directly at RT for 30min), then were blocked with 5% goat serum in PBST for 30 min. Primary antibody incubation was performed at RT for 1-2 h. Sections were washed with PBST, then secondary antibody incubation was performed at RT for 1 h. Hoechst 33342 (Thermo Fisher Scientific) was diluted 1:5000 in PBS and applied for 12 minutes for staining nuclei. Retinal explant sections and flatmounts were imaged using an Olympus FV1200 confocal microscope. Antibodies used: Rabbit anti-GFP (Thermo Fisher Scientific, A11122, 1:1000), Goat anti-Rabbit IgG (H+L) Highly Cross-Adsorbed Secondary Antibody Alexa Fluor Plus 488 (Thermo Fisher Scientific, A32731, 1:1000), PNA (Thermo Fisher Scientific, L32460, 1:200).

## Acknowledgments

We are deeply grateful to the human donors and their families who made this study possible. We thank the Center for Organ Recovery and Education for facilitating the use of human donor eyes. Animal husbandry was performed by University of Pittsburgh Division of Laboratory Animal Resources (DLAR). Deep sequencing was performed at UPMC Genome Center. This work used the Extreme Science and Engineering Discovery Environment (XSEDE),^37^ which is supported by National Science Foundation grant number ACI-1548562. Specifically, it used the Bridges-2 system, which is supported by NSF award number ACI-1445606 and ACI-1928147, at the Pittsburgh Supercomputing Center (PSC). This research was also supported in part by the University of Pittsburgh Center for Research Computing through the resources provided. We appreciate help from Dr. Yuanyuan Chen and Dr. Abhishek Vats in establishing the retinal explant culture protocol. We acknowledge support from NIH CORE Grant P30 EY08098 to the Department of Ophthalmology, from the Eye and Ear Foundation of Pittsburgh, and from an unrestricted grant from Research to Prevent Blindness, New York, NY. Funding was provided by the UPMC Immune Transplant and Therapy Center, Foundation Fighting Blindness, the Research to Prevent Blindness (LB; Career Development Award), and through a scholarship from the China Scholarship Council and research funding from The Third Xiangya Hospital and Xiangya School of Medicine to ZX.

## Author contributions

ZX: Conceived, planned and executed experiments. Analyzed data. Wrote the manuscript. BEO: Conceived, planned and executed experiments. Analyzed data. Wrote the manuscript. MEJ: Conceived, planned and executed experiments, analyzed data. Wrote the manuscript. LCB: Supervised work. Conceived, planned and executed experiments. Analyzed data. Wrote the manuscript.

## Data availability

Data is available upon request.

## Supplementary Materials

**Figure S1.**
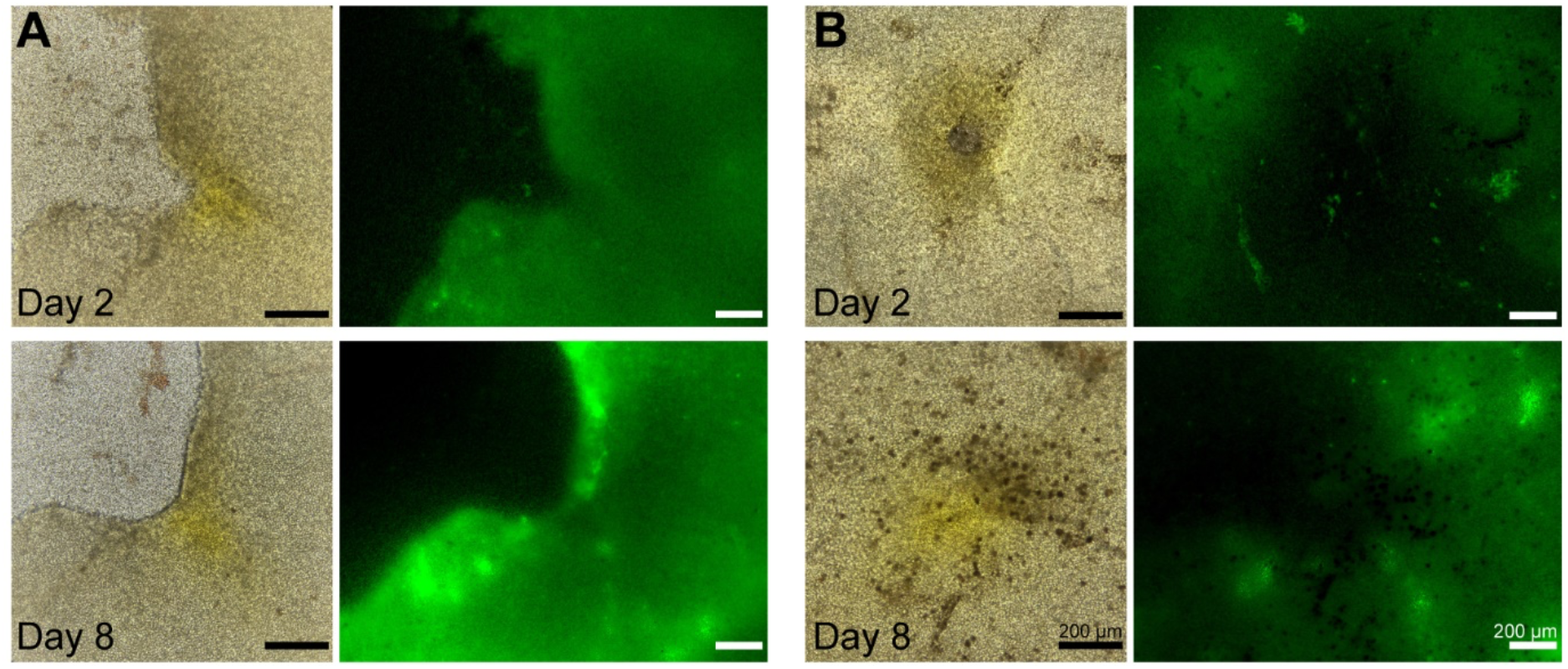
GFP expression in the foveal pit of retinal explants. (A) Foveal pit located on the edge of the explant. The macula can be identified as the area with yellow pigmentation. (B) Foveal pit in the center of the explant. Bright field images are on the left panel, and FITC images showing GFP are on the right panel of each figure. GFP expression started from the edge of the explant with growing intensity and area towards the center of the explant over time.

**Figure S2.**
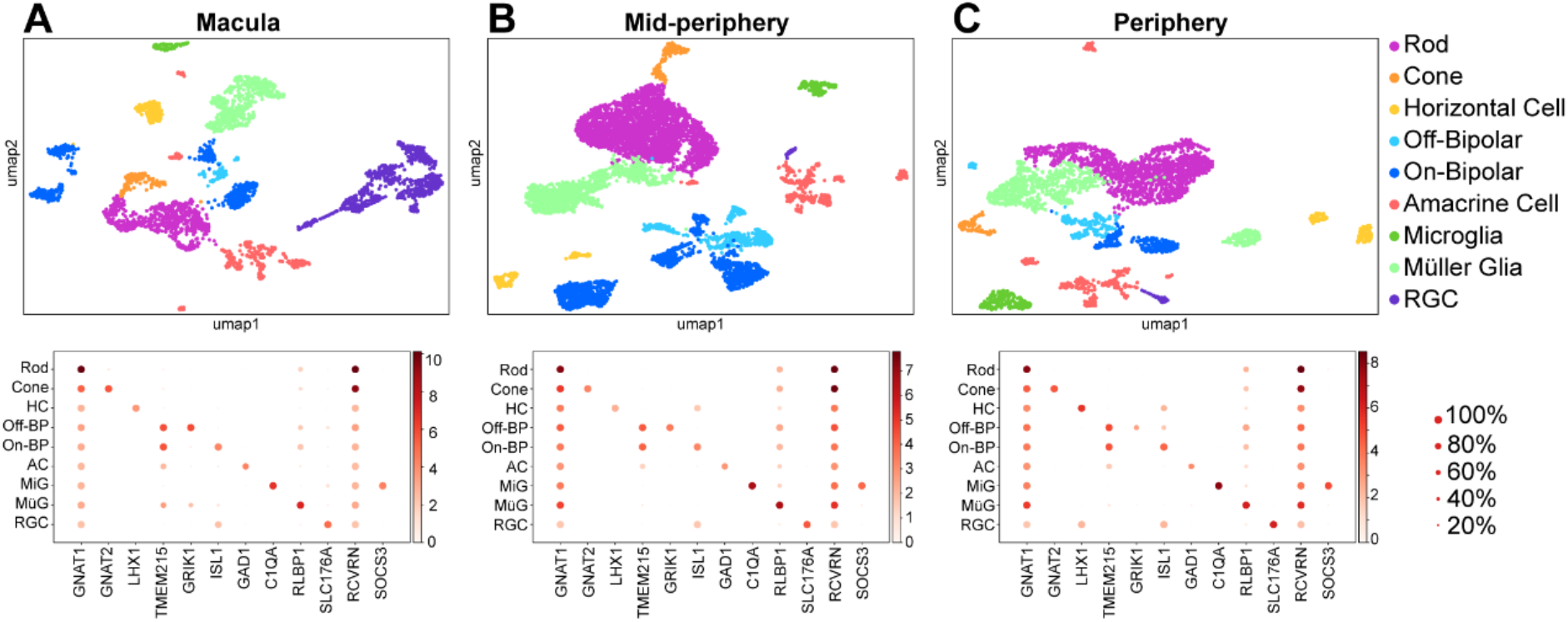
Cell clustering and identification of fresh human retina. Major retinal cell types are clustered and identified from macular, mid-peripheral, and peripheral retinal samples (2 samples from left and right eye per anatomical location) of the fresh retina from donor#2.

**Table S1.**
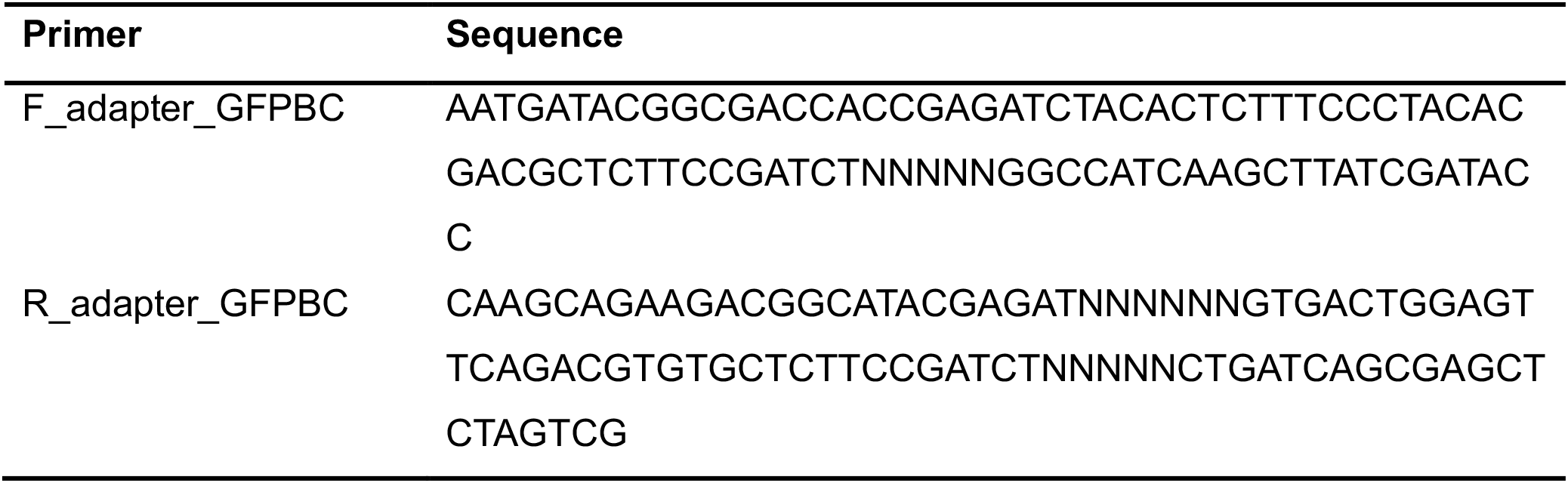

